# Assassination Tango: An NLR/NLR-ID immune receptors pair of rapeseed co-operates inside the nucleus to activate cell death

**DOI:** 10.1101/2021.10.29.466428

**Authors:** Glykeria Mermigka, Argyro Amartolou, Adriani Mentzelopoulou, Niki Astropekaki, Panagiotis F. Sarris

## Abstract

Plant immunity relies on cell-surface receptors and intracellular NLR immune receptors. Some plant NLRs carry integrated domains (IDs) that that mimic authentic pathogens effector targets. We report here the identification in *Brassica napus* of the genetically linked NLR/NLR-ID pair, *Bn*RPR1 and *Bn*RPR2. The NLR-ID carries two ID fusions and their mode of action conforms to the proposed “integrated sensor/decoy” model. The two NLRs interact and the heterocomplex localizes in the plant-cell nucleus and nucleolus. However, the *Bn*RPRs pair does not operate through a negative regulation as it was previously reported for other NLR-IDs. Cell death is induced only upon co-expression of the two proteins and it is dependent on the helper genes EDS1 and NRG1. Truncations of the IDs of *Bn*RPR1 results in cytoplasmic localization and compromises cell death activation. Expression, using the native promoter in *Nicotiana* species, led to a conditional cell death induction. However, this was not the case for the transgenic Arabidopsis, where no cell death was observed. In summary, we describe a new pair of NLR-IDs with interesting features in relation to its regulation and the cell death activation.

## Introduction

In order to protect themselves from pathogenic microorganisms, plants deploy a versatile defense toolkit/network. This extends from mechanical barriers (e.g. cell walls) to sophisticated biomolecular sensors such as specialized membrane receptors and various Resistance (R) proteins. R proteins could be intracellular immune receptors that recognize microbial-secreted pathogenicity components, known as effector proteins, and initiate a chain of events leading to immune response (<u>E</u>ffector <u>T</u>riggered Immunity, ETI). ETI often drives to a programmed cell death in order to restrict pathogens growth and spread (Dangl and Jones, 2001). Most of the known R proteins belong to the NB-LRR family since they contain a nucleotide binding domain (NB) followed by leucine rich repeat domain (LRR), while their amino-(N-) and carboxyl-(C-) terminal domains may vary. Depending on their amino terminal part, they are divided into three major groups; a) the TIR-NBS-LRR (TNL) containing a TIR (Toll/interleukin1 receptor) domain,; b) the CNLs containing a Coiled-Coil (CC) domain and c) the RPW8-NBS-LRR (RNLs) that harbor a RPW8 (RESISTANCE TO POWDERY MILDEW 8) N-terminal domain (Meyers et al., 1999; Pan et al., 2000; Xiao et al., 2001; Meyers et al., 2003).

Some NLRs have additional non-canonical domains which enable perception of pathogen effectors. Studies reveal different NLRs of *Arabidopsis thaliana* and rice that have such domains, termed Integrated Domains (IDs) (Cesari et al., 2014a; Le Roux et al., 2015; Maqbool et al., 2015; Sarris et al., 2015). The so called “integrated sensor/decoy” (ISD) model has been proposed to accommodate their mode of function (Cesari et al., 2014a; Sarris et al., 2015; Wu et al., 2015). According to this model, IDs serve as decoys for effectors, by mimicking their real subcellular target in the host cell. When interacting with the effector, potential conformational changes are induced to one NLR of the pair, which in turn are sensed from the partner NLR, which is usually genetically and physically linked (Le Roux et al., 2015; Maqbool et al., 2015; Sarris et al., 2015).

One such pair of NLRs revealing the above features and mode of action is the Arabidopsis TNL pair RRS1/RPS4. RRS1 (Resistance to *Ralstonia solanacearum* 1) contains an ID that reveals homology to the WRKY family of plant transcription factors which mimics the effector target of the host. Sarris *et al*. (Sarris et al., 2015) and Le Roux *et al*. (Le Roux et al., 2015), showed independently that PopP2, an acetyltransferase effector of *Ralstonia solanacearum*, binds and acetylates the WRKY domain of RRS1 thus activating immunity. Similarly, the unknown function effector AvrRps4, of *Pseudomonas syringae*, targets the WRKY domain of RRS1 activating defense (Le Roux et al., 2015; Sarris et al., 2015). The genetically linked NLR protein RPS4 (Resistance to *Pseudomonas syringae* 4), which physically interacts with RRS1 and forms heterocomplex, “senses” the conformational change of RRS1 and initiates the immune response (Sohn et al., 2014; Huh et al., 2017). Interestingly, the cleavage of the WRKY domain from RRS1 protein, leads to constitutive defense activation that is RPS4 dependent. This indicates a self-control role for this WRKY ID, in order to keep the RRS1/RPS4 system inactive in an elicitors’ free environment (Sarris et al., 2015; Huh et al., 2017).

Similarly, the CNL pair RGA4/RGA5 acts cooperatively for the recognition of two unrelated effectors of the rice blast fungus *Magnaporthe oryzae*, AVR-PiA and AVR1-CO39 (Okuyama et al., 2011; Cesari et al., 2013). In the inactive state (absence of effector) RGA5 forms heterocomplexes with RGA4 thus repressing the onset of the cell death signaling pathway, mediated by the executor RGA4. In the presence of *M. oryzae*, the C-terminal part of RGA5 harboring and ID “Heavy Metal Association” (HMA) domain, physically associates with pathogens effector protein(s) leading to conformational changes of RGA5 ultimately leading to RGA4 derepression (Cesari et al., 2014b). An analogous mode of action has been reported for the rice CNL Pik-1/Pik-2 pair. (Ashikawa et al., 2008; Maqbool et al., 2015). The HMA domain of Pik-1 also functions as a decoy for the recognition of another effector of *M. oryzae*, the AVR-Pik. *In silico* analyses has shown that a large number of NLRs harbor IDs and that some of these IDs overlap with previously identified pathogen targets (Kroj et al., 2016; Sarris et al., 2016; Bailey et al., 2018).

Despite the importance of NLRs in plant defense, regulating their expression is crucial for maintaining a balance between normal plant growth and adequate defense. Overexpression or ectopic expression of NLRs might lead to unwanted activation of immunity or developmental problems while inadequate expression might lead to susceptibility against specific pathogens (Yang and Hua, 2004; Zhang et al., 2004; Kim et al., 2010; Li et al., 2010; Palma et al., 2010; Cesari et al., 2013; Chae et al., 2014). Thus, NLR expression is a tightly regulated process occurring at various levels, from the time of their transcription until the time of their protein degradation (Li et al., 2015; Borrelli et al., 2018; Lai and Eulgem, 2018).

Here, we report the identification of two TNLs in the genome of *Brassica napus* that are transcribed in a head-to-head orientation. According to our data, these TNLs adopt the “integrated sensor/ decoy” (IDS) mode of action and lead to a cell death activation when ectopically expressed while their expression seems to be regulated at the post-transcriptional level.

## Results

### Identification and characterization of the *BnRPR1/BnRPR2* NLR pair from *B. napus*

In order to identify NLRs with integrated domains fitting into ISD model we used Arabidopsis RRS1 as the query sequence in our blast search against the genome of the crop plant *Brassica napus* (cultivar ZS11) and filtered the results based on the presence of a second genetically linked putative NLR. We chose to study a pair of putative NLRs which we termed *Brassica napus RESISTANCE PAIRED RECEPTOR 1 & 2* (*BnRPR1 & BnRPR2*). These genes contain some interesting features: a) they are two NLRs genetically linked with a head-to-head orientation, which potentially implies a combined mode of action; b) *BnRPR1* codes for a NLR that contains two integrated domains (IDs) at the N- and C-terminal end of the protein. The N-terminal domain, B3, has been found in five major groups of plant transcription factors (Romanel et al., 2009). The C-terminal domain, TFSIIN, is a part of the transcription elongation factor IIS (TFSII) (Wind and Reines, 2000). The presence of these IDs in an NLR protein suggest a possible mode of action according to the ISD-model; c) part of their mRNA 5’ ends, overlap, potentially suggesting an involvement of a post-transcriptional regulation mechanism. The above mentioned features are summarized in Figure 1A.

**Figure 1:**
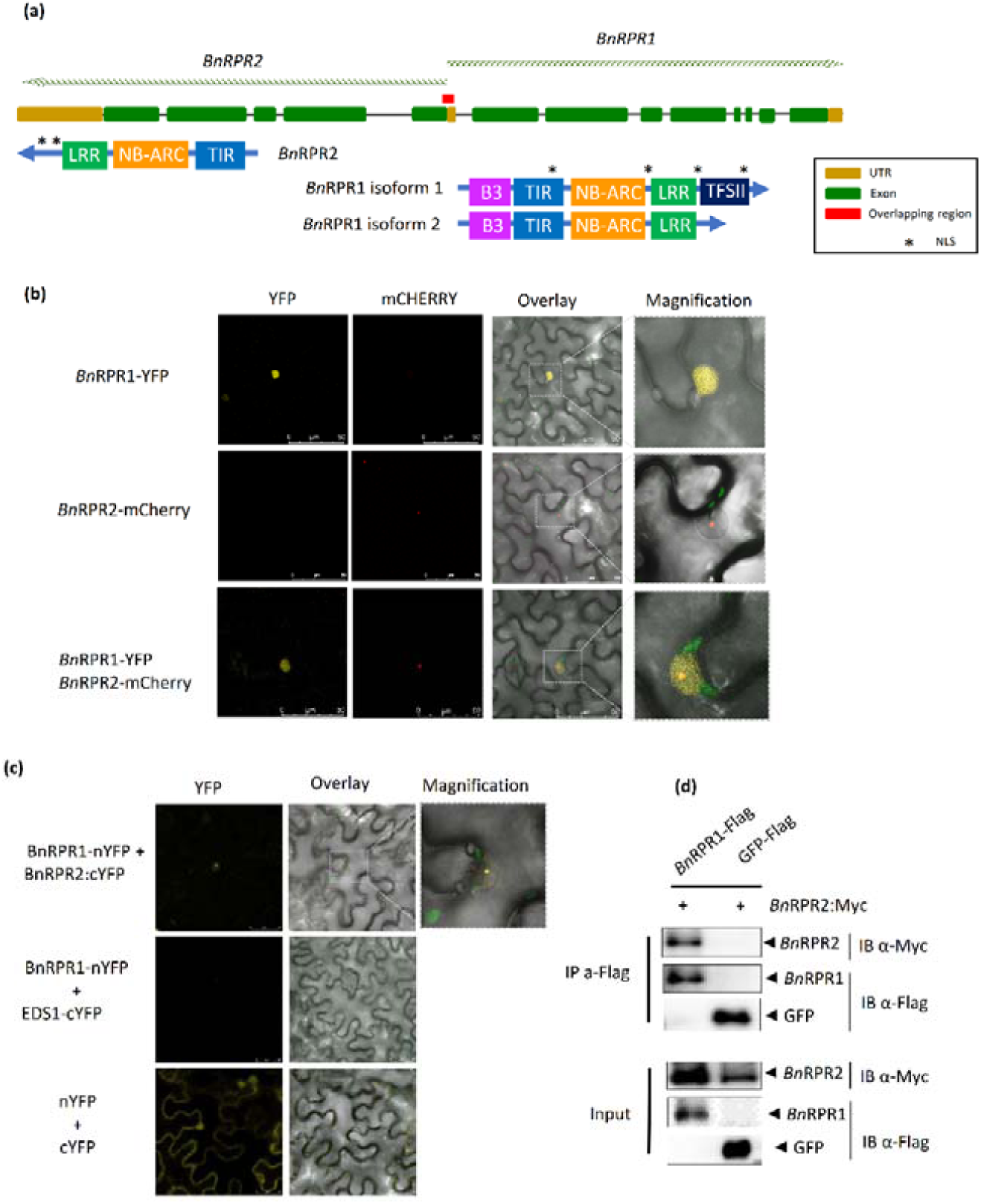
Characterization of the genetically linked NLRs, BnRPR1 and BnRPR2. (a) Graphical representation of the *BnRPR* genetic locus and their corresponding proteins. BnRPR1 and BnRPR2 are genetically linked and transcribed in a head- to-head orientation with an overlapping region that corresponds to a small part of their transcripts. According to *in silico* analysis, BnRPR1 codes for two isoforms, with the second lacking the TFIISN domain. Asterisks in each protein mark the nucleus localization signal, as predicted by LOCALIZER (localizer.csiro.au). (b) Subcellular localization of *Bn*RPR1 and *Bn*RPR2. *Bn*RPR1 shows a diffused pattern inside the nucleus while BnRPR2 is localized in certain foci inside the nucleus. Leaf sections of *N. benthamiana* plants that transiently express 35S:*BnRPR1-YFP* or 35S:*BnRPR2-m*Cherry were analyzed with confocal microscopy. (c) BiFC assays reveal close proximity of *Bn*RPR1 and *Bn*RPR2. A signal could be observed in certain foci inside the nucleus. Leaf sections *N. benthamiana* plants transiently co-expressing 35S:*BnRPR1-nYFP* and 35S:*BnRPR2-cYFP* were analyzed with confocal microscopy. Negative controls (35S:BnRPR1-nYFP/35S:EDS1-cYFP and 35S:nYFP/35S:cYFP) were also included. (d) Co-IP assay showing the interaction between *Bn*RPR1 and *Bn*RPR2. 35S:*BnRPR1-Flag* and 35S:*BnRPR2-Myc* were transiently co-expressed in *N. benthamiana leaves*. The protein lysate was immunoprecipitated with α-Flag. *Bn*RPR2 immunoprecipitated with *Bn*RPR1 while in the negative control used (35S:GFP-Flag/35S:*BnRPR2-Myc*) no signal could be observed. IB= Immunoblot

Initially, we confirmed the expression of *BnRPR1* and *BnRPR2* in the leaves of 10 days old *B. napus* cv. ZS11 plants. Our RT-PCR results showed both genes are expressed at the examined tissues (Sup. Figure 1). Next, in order to study the subcellular localization of both *BnRPR1* and *BnRPR2*, we cloned the genomic sequence of each gene into plant expression vectors. The genes were tagged at the C-terminus with an appropriate fluorophore and their expression was driven by the 35S promoter of Cauliflower mosaic virus (CaMV). Using agroinfiltration we expressed both constructs (35S:*BnRPR1-YFP*, 35S:*BnRPR2-mCherry*) in *Nicotiana benthamiana* leaves and their localization was examined using confocal microscopy, two days post agroinfiltration (dpa) (Figure 1B). When expressed separately, *Bn*RPR1 and *Bn*RPR2 localized in the nucleus, however in different patterns. *Bn*RPR1 showed a diffused pattern inside the nucleus while *Bn*RPR2 localized in certain foci inside the nucleus, which resemble nucleolus. Upon co-expression, their localization remained as above.

Since genetically linked NLRs tend to cooperate (Duxbury et al., 2016), we used Bimolecular fluorescence complementation (BiFC) and co-immunoprecipitation (co-IP) assays to test if the same applies to the BnRPRs (Figure 1C & 1D). *BnRPR1-nYFP* was transiently co-expressed with *BnRPR2-cYFP* in *N. benthamiana* leaves and leaf sections were visualized using confocal microscopy. Our BiFC assay revealed a signal inside the nucleus pointing to a nuclear association of *Bn*RPR1 and *Bn*RPR2 (Figure 1C). This is consistence with our previous co-localization studies. The association of *Bn*RPR1 and *Bn*RPR2 was also confirmed using co-IP assays that were performed by co-expressing *BnRPR1-FLAG* and *BnRPR2-MYC* in *N. benthamiana* leaves (Figure 1D). Overall, our data indicate that *Bn*RPR1 and *Bn*RPR2 are two NLRs that interact inside the plant-cell nucleus.

### Cell death induction due to BnRPR1 and BnRPR2 co-expression is EDS1 and NRG1 dependent

During our co-agroinfiltration assays we observed that co-expression of *BnRPR1-YFP* and *BnRPR2-m*Cherry in *N. benthamiana*, as well as, in *N. tabaccum*, leads to induction of cell death (Figure 2A). The same effect was not detected when the two genes were separately expressed. In order to investigate whether the tags used might interfere with the correct protein folding thus leading to cell death activation, we repeated the assay with the NLRs tagged with different epitopes or without epitopes (Sup. Figure 2). The cell death activation was induced in all cases, although in one combination (*BnRPR1-nVENUS*/*BnRPR2-cCFP*) it was slight delayed. Next, we investigated whether this cell death activation requires the activity of two helper NLRs, the N REQUIRED GENE 1 (NRG1) (Peart et al., 2005) and the ENHANCED DISEASE SUSCEPTIBILITY 1 (EDS1) (Dongus and Parker, 2021). For this we performed the assays on *eds1* and *nrg1 N. benthamiana* knock-out lines (Ordon et al., 2017; Qi et al., 2018) (Figure 2B). The co-expression of *BnRPR1-YFP*/*BnRPR2-m*Cherry lead to the induction of a cell death only in w.t. *N. benthamiana* plants. Our data reveal that the cell death immune response triggered by the ectopic co-expression of *Bn*RPR1 and *Bn*RPR2 requires functional EDS1 and NRG1 proteins.

**Figure 2:**
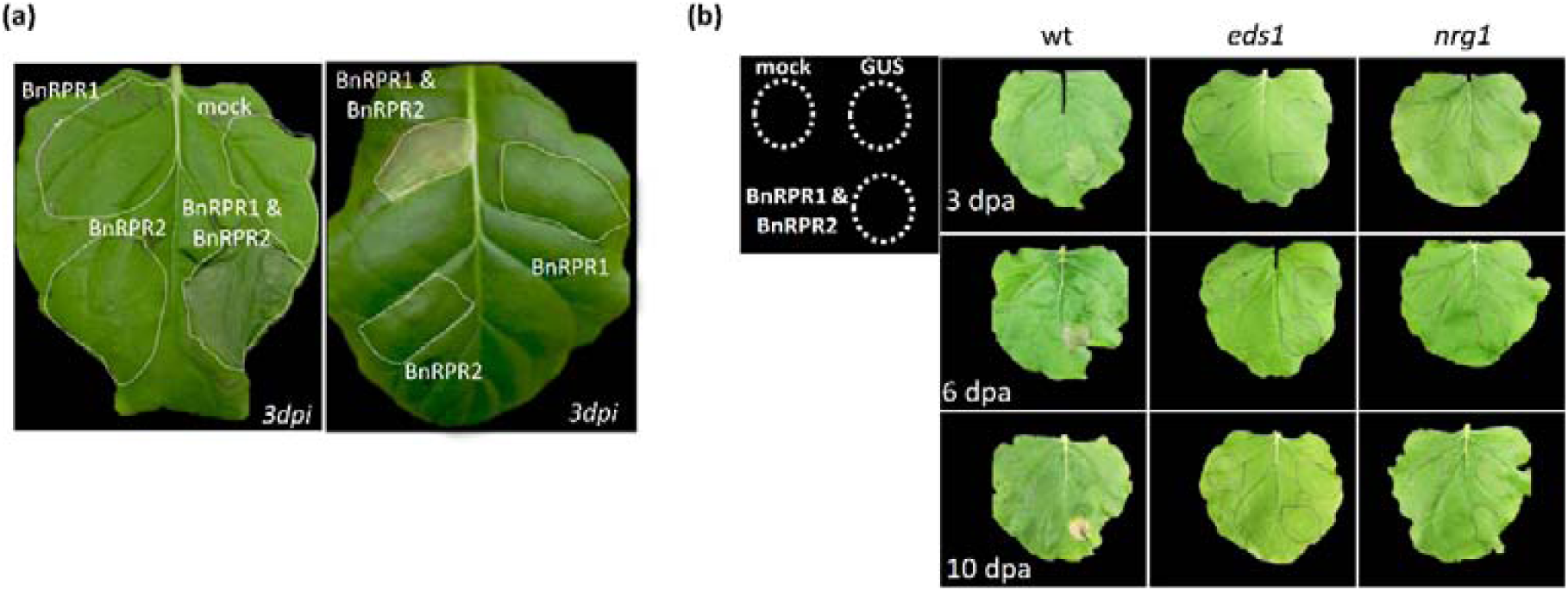
Transient expression of *Bn*RPR1 and *BnRPR2* in *Nicotiana* species triggers cell death which is EDS1 and NRG1 dependent. (a) Transient co-expression of *Bn*RPR1 and *Bn*RPR2 *N. benthamiana* (left panel) and *N. tabacum* (right panel) leads to cell death. 35S:*BnRPR1-YFP* and 35S:*BnRPR2-m*Cherry were transiently expressed in *Nicotiana* leaves separately or together. Three days post agro-infiltration (dpi) cell death could be observed. (b) The cell death induced by *BnRPR1* and *BnRPR2* is EDS1 and NRG1 dependent. *BnRPR1* and *BnRPR2* were transiently co-expressed in leaves of *wt*, eds1 and nrg1 *N. benthamiana* plants and the progression cell death was monitored during a 10 day period. Transient co-expression of *BnRPR1* and *BnRPR2* leads to cell death only in w.t. plants. As negative controls agroinfiltration with GUS and mock infiltration were included.

### Changes in the subcellular localization of BnRPRs impacts on the onset of cell death

*Bn*RPR1 contains two integrated domains (IDs); a B3 domain at the N-terminus and a TFSIIN domain at the C-terminus of the protein. In order to investigate their role in defense activation, we generated three truncated versions of *Bn*RPR1: a) lacking the B3 domain (BnRPR1*Δ*B3), b) lacking the TFSIIN domain (*Bn*RPR1*Δ*TFSIIN) which corresponds to the second isoform of the protein (see above) and c) lacking both ID domains (*Bn*RPR1*Δ*B3*Δ*TSSIIN). All these truncations were tagged with a fluorophore and expressed under the 35S promoter. The constructs were transiently expressed by agroinfiltration in *N. benthamiana* leaves and their localization was examined by confocal microscopy two dpa, in the absence or presence of *Bn*RPR2 (Figure 3 & Sup. Figure 3). When co-expressed with *Bn*RPR2, the truncated proteins lacking the TFSIIN domain reveal higher levels of localization in the cytoplasm, a pattern that was also observed when the truncated proteins were expressed alone (Figure 3B & Sup. Figure 3B). Unexpectedly, a difference in the subcellular localization of *Bn*RPR2 when expressed with the truncated versions of *Bn*RPR1 was also detected. In more details, when *Bn*RPR2 was co-expressed with the w.t. version of *Bn*RPR1 a signal was detected only in the nucleolus, as expected from the experiments mentioned above (Fig. 1B). Upon co-expression with the truncated versions of *Bn*RPR1 lacking the TFSIIN domain, *Bn*RPR2 was also detected in the cytoplasm (Figure 3B).

**Figure 3:**
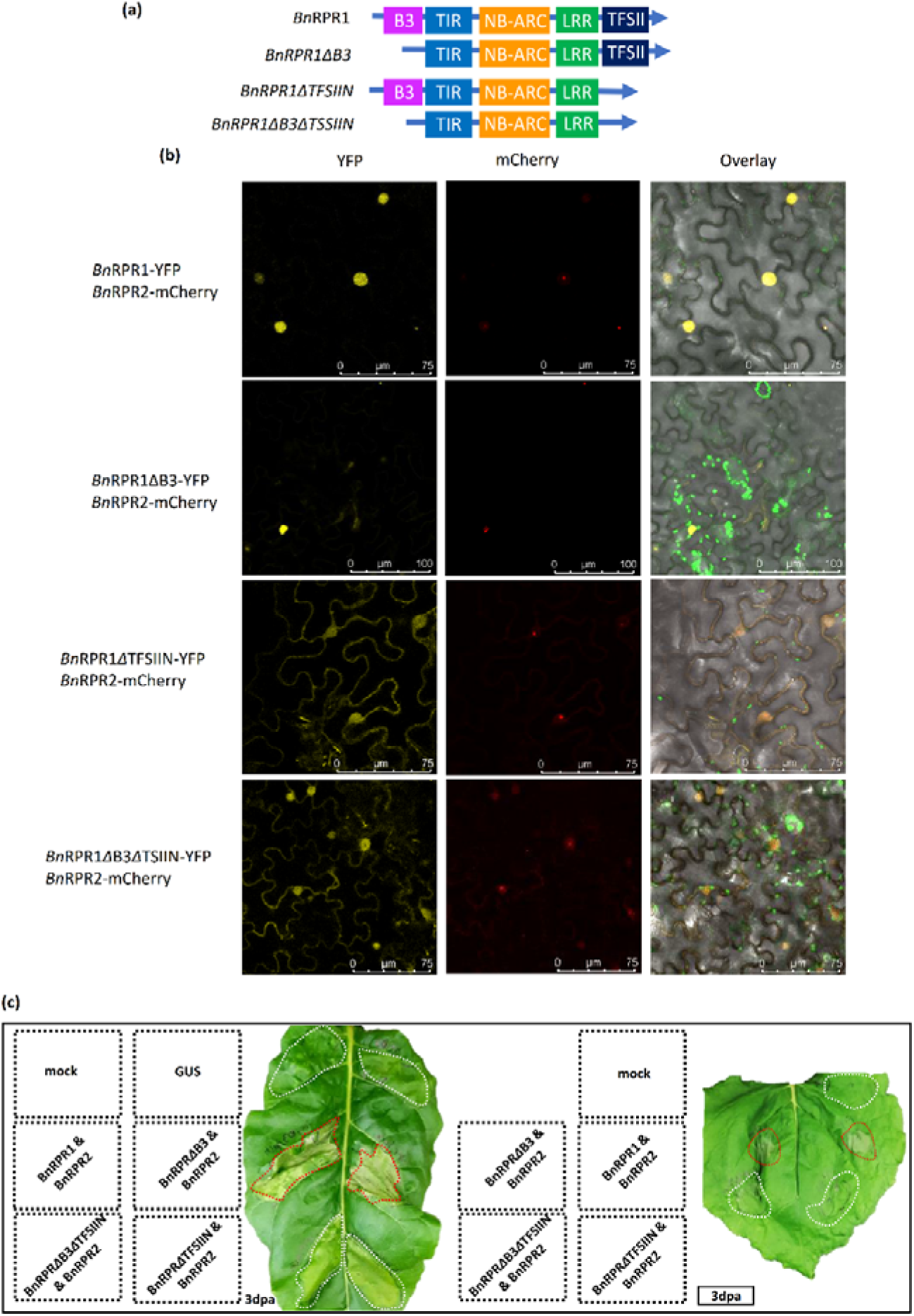
Subcellular localization and cell death induction of the three *Bn*RPR1 truncations. (a) Graphical representation of the different truncations of *Bn*RPR1, lacking the B3 domain (*Bn*RPR1ΔB3), the TFSIIN domain (*Bn*RPR1ΔTFSIIN) or both (BnRPR1ΔB3ΔTFSIIN). (b) Truncations of BnRPR1 are also identified in the cytoplasm while BnRPR2 can also be identified in the cytoplasm when co-expressed with these truncations. BnRPR2-mCherry was transiently co-expressed with *Bn*RPR1-YFP and the three truncations (*BnRPR1ΔB3-YFP, BnRPR1ΔTFIIN-YFP* and *BnRPR1ΔB3ΔTFIIN-YFP* in *N. benthamiana* leaves and the leaf sections were analyzed with confocal microscopy two dpa. Truncations of *BnRPR1* have a nucleocytoplasmic localization a pattern also observed for *Bn*RPR2 only when co-expressed with the truncated versions of *Bn*RPR1 lacking the TFSIIN domain. (c) The truncated versions of *Bn*RPR1 lacking the TFSIIN domain show a delayed cell death. *Bn*RPR2-mCherry was transiently co-expressed with BnRPR1-YFP and the three truncations (BnRPR1ΔB3-YFP, BnRPR1ΔTFIIN-YFP and *Bn*RPR1ΔB3ΔTFIIN-YFP in *N. tabaccum* leaves and cell death was monitored. Three dpa cell death could be observed in the BnRPR1-YFP and BnRPR1ΔB3-YFP. Red dots denote the macroscopic HR symptom.

The truncated versions of *Bn*RPR1 were also used to study their effect on the activation of the cell death. The w.t. and the three truncated versions of *Bn*RPR1 were co-expressed with *Bn*RPR2 in *N. benthamiana* and *N. tabaccum* leaves, using agroinfiltration and the onset of cell death induction was monitored (Figure 3C & Sup. Figure 3C). Three days post agroinfiltration (3dpa) the cell death induction was activated where *BnRPR2* was co-expressed with *BnRPR1* and *BnRPR1ΔB3* but not when *BnRPR2* was co-expressed with *Bn*RPR1*Δ*TFSIIN or *Bn*RPR1*Δ*B3*Δ*TSSIIN (Figure 3C & Sup. Figure 3C).

Our data indicate that the truncated versions of *Bn*RPR1 lacking the TFSIIN domain show a different subcellular localization compared to w.t. *Bn*RPR1 protein and the *Bn*RPR1*Δ*B3 truncation, since they could also be detected in the cytoplasm. Additionally, their co-expression with *Bn*RPR2 has also an impact in the localization of *Bn*RPR2, since in their presence *Bn*RPR2 adopts a cytoplasmic pattern in addition to its nuclear localization. Finally, regarding the effect of the *Bn*RPR1 truncations on the onset of the cell death activation, we observed that the changes in the subcellular localization leads to differences in the activation of cell death.

### Co-expression of BnRPR1 and BnRPR2 leads to decreased transcript and protein levels

Our confocal analysis revealed a signal intensity reduction upon w.t. *Bn*RPR1/*Bn*RPR2 co-expression. In order to evaluate this observation, we measured the transcript and protein levels of both *BnRPR1* and *BnRPR2* (Figure 4). Both the *35S:BnRPR1-FLAG* and *35S:BnRPR2-MYC* constructs were transiently expressed together in *N. benthamiana* leaves or separately with the *35S:GUS-MYC* construct. Transcripts and proteins levels from each treatment were validated using a reverse transcription quantitative PCR (RT-qPCR) and western blot, respectively (Figure 4). The transcript levels of *Bn*RPR1 and *Bn*RPR2 were downregulated upon co-expression (Figure 4B), which resulted in reduced protein levels (Figure 4C). Overall, our data reveal that co-expression of *Bn*RPR1 and *Bn*RPR2 leads to reduced transcript and protein levels.

**Figure 4:**
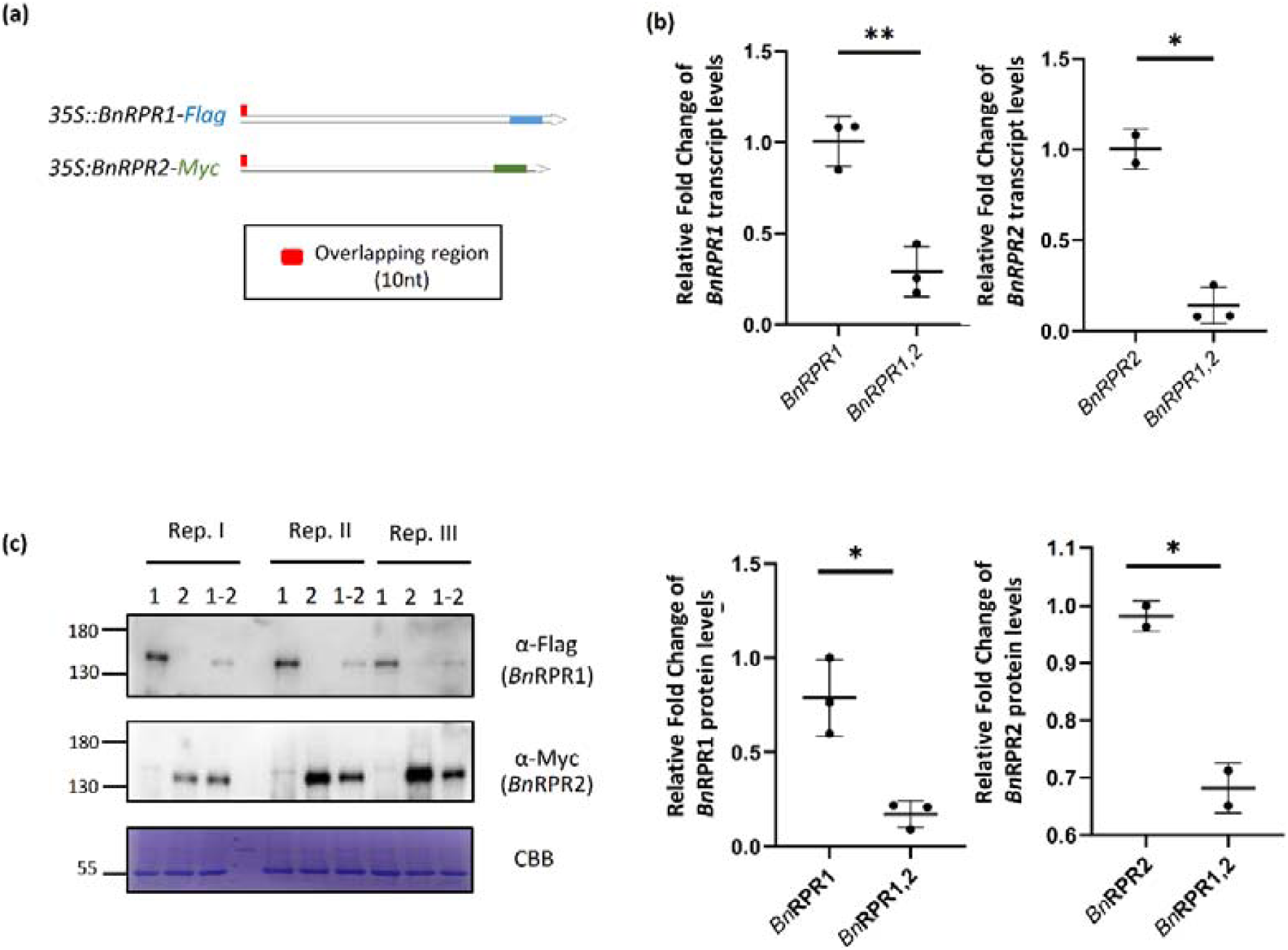
BnRPR1 and BnRPR2 are downregulated and transcript and protein level when co-expressed. (a) Graphical representation of the *BnRPR* constructs used. Red rectangle depicts the overlapping region of the two *BnRPR* transcripts (b) RT-qPCR analysis showing the downregulation of *Bn*RPR1 & 2 when they are co-expressed. *Bn*RPR1 and *Bn*RPR2 were transiently expressed in *N. benthamiana* leaves either alone (with *35S:GUS*-*MYC*) or together. cDNA generated from DNaseI treated RNA extracted from the leaves was used for qPCR analysis. The expression level of each gene was normalized to the expression of the housekeeping gene UBI and NPTII. Bars represent mean of SD (n= 2-3 biological replicates). Bars marked with asterisks indicate significant differences (Student’s t-test), * P-value < 0.05, ** P-value < 0.005). (c) Western blot analysis showing the downregulation of *Bn*RPR1 & 2 when they are co-expressed. The same samples used for RT-qPCR analysis were used to perform western blot analysis. The same membrane was initially analyzed with a-Flag and after stripping the a-Myc antibody was used. Signal quantification was performed with Azzure spot 2.0 (AzureBiosystems). Bars represent mean of SD (n= 2-3 biological replicates). Bars marked with asterisks indicate significant differences (Student’s t-test), * P-value < 0.05.

### Comparison of BnRPRs transcript levels and the cell death induction in different genetic backgrounds and promoter sequences

The cell death induction that was observed when *BnRPR1* and *BnRPR2* were ectopically expressed, might be the outcome of overexpression or/and ectopic expression, a case frequently reported for NLRs (Yang and Hua, 2004; Zhang et al., 2004; Kim et al., 2010; Li et al., 2010; Palma et al., 2010; Cesari et al., 2013; Chae et al., 2014). To investigate this, we used the 35S promoter constructs, as well as, the native promoter of the *Bn*RPRs. The 35S promoter is generally considered as a strong promoter (Odell et al., 1985), leading to high expression levels of downstream sequences, that could be responsible of the cell death phenotype. In order to evaluate the effect on the cell death activation when the *Bn*RPRs have expression levels similar to the ones expressed in *B. napus*, we cloned into a plant expression vector the gDNA of the entire *BnRPRs* genetic locus, containing the native promoter and the 3’ and 5’ UTR sequences. This construct was used for agroinfiltration assays in *Nicotiana* species, as well as, for the generation of transgenic *A. thaliana* (Col-0) lines that express the *BnRPRs* locus (Col^BNRPR^).

Transient expression of the *BnRPRs* locus in *N. tabaccum* leaves led to a cell death activation, however, in a small number of the examined plants (3 out of 10 plants in independent experiments). Since our plants were initially grown in greenhouse conditions with temperature fluctuations, we decided to repeat these experiments in controlled conditions in a side-by-side manner using two groups of plants. The two groups were agroinfiltrated with the *BnRPRs* locus and then placed at 26 °C and 22°C, respectively. Both plant groups were monitored for cell death induction. In these experiments, the co-expression of *35S:BnRPR1* and *35S:BnRPR2* was used as a positive control. We observed that 6/6 plants (in three independent experiments) grown at 26 °C, reveal a strong cell death phenotype. In contrast to the plants that were grown at 22 °C, which did not develop cell death (Figure 5A). The cell death observed in 26 °C upon *BnRPRs* locus expression, showed a 24 hrs delay compared to cell death induction upon the *35S:BnRPR1* and *35S:BnRPR2* co-expression. However and concerning our transgenic Arabidopsis Col^BNRPR^ lines, they were viable with developmental phenotypes identical to w.t. plants (Figure 5B and Sup. Figure 4).

**Figure 5:**
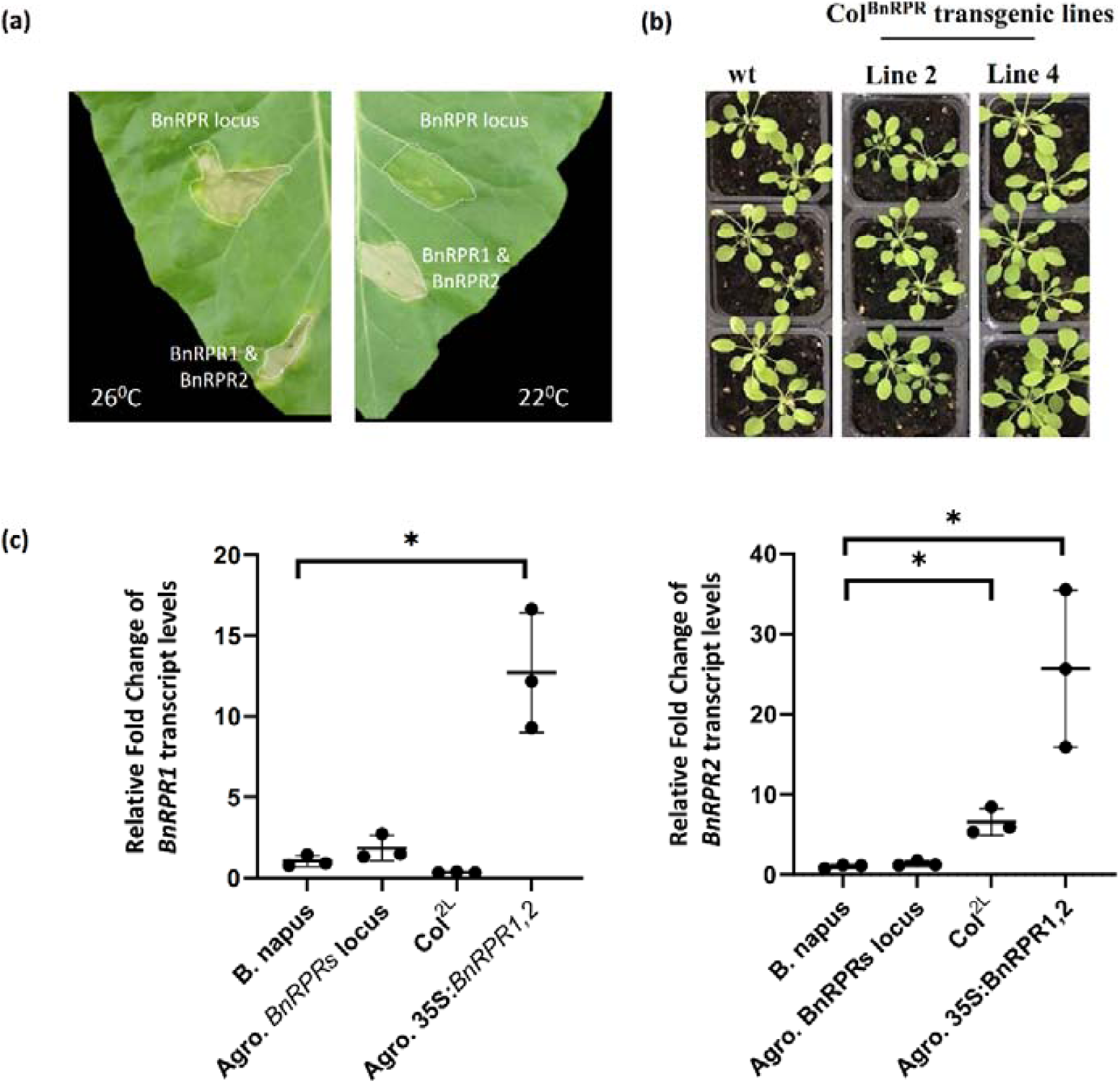
Effect of *BnRPR1* and *BnRPR2* expression on the cell death and their expression levels in the different backgrounds. (a) Transient ectopic expression of the *BnRPR* locus leads to cell death conditionally. *Bn*RPR locus was transiently expressed in leaves of *N. tabaccum* plants grown in different conditions (Cond. I: 22 ° C 50% RH vs Cond. II: 26 ° C 80% RH). As a positive control for HR induction 35S:*BnRPR1-YFP* and 35S:*BnRPR-m*Cherry were co-expressed in the same leaves. In both conditions the 35S:*BnRPR1-YFP/35S:BnRPR2-m*Cherry expression lead to cell death, while cell death was only detected at Cond. II when *Bn*RPR locus was expressed with one day delay. The experiment was repeated twice in a total of six plants with similar results. (b) *A. thaliana* transgenic lines constitutively expressing the *BnRPR* locus (Col^BNRPR^) have a phenotype similar to w.t. plants. T2 generation *A. thaliana* Col^BNRPR^ transgenic lines are compared to w.t. plants. (c) Accumulation levels of *BnRPR1* and *BnRPR2* in the different genetic backgrounds. Real-time PCR was conducted in cDNA from: i) *N. benthamiana* leaves transiently expressing 35S:*BnRPR1-YFP* and 35S:*BnRPR2-m*Cherry, ii) *N. benthamiana* leaves transiently expressing the *BnRPR* locus, iii) A. thaliana leaves constitutively expressing the *BnRPR* locus (Col^2L^) and iv) *B. napus leaves*. The highest levels were observed in condition (i). In all cases *ACTIN* was used as a reference gene (n=3, p < 0.05).

Using RT-qPCR we next examined the expression levels of *BnRPR1* and *BnRPR2* in *B. napus* leaves, in *N. benthamiana* leaves agroinfiltrated with the *BnRPRs* locus or the combination of 35S:BnRPR1/35S:BnRPR2 and in the transgenic Arabidopsis line 2 (Col^2L^) expressing *BnRPRs* (Figure 5C). In all cases actin was used as the reference gene. In their native environment the tested genes are expressed at low levels, while when transiently expressed their expression was elevated by 10 to 20 fold changes, compared to the levels of *BnRPR1* and *BnRPR2* detected in *B. napus* (Figure 5C). Altogether, the native promoters of the two *Bn*RPR NLRs lead to low expression levels but when the locus is ectopically expressed in tobacco it leads to a conditional onset of cell death activation.

## Discussion

The discovery of new NLRs has been the focus of various plant pathology laboratories and plant breeding programs around the world, since they represent a sustainable means to control plant diseases. However, by investigating their intrinsic features, we can also enrich our knowledge on the mode of action and regulation of plant defense mechanisms. A special class of NLRs that has emerged the last decade concerns the NLR proteins that harbor integrated domains and usually co-operate with other genetically linked NLRs (Grund et al., 2019).

Focus of the present study is the identification and characterization of a pair of genomically linked NLRs, which are encoded in the genome of *B. napus* (var. ZS11), the *Bn*RPR1 and *Bn*RPR2. Our data reveal that these two NLRs are functional since upon co-expression they induce cell death in *Nicotiana* species (Figure 2A & Sup. Figure 2) and it seems to fit the “integrated sensor/decoy” (ISD) model. *Bn*RPR1 contains two IDs (Figure 1A) and it physically interacts with *Bn*RPR2. This interaction takes place in the nucleus, with the nucleolus being the most prominent site of complex formation upon co-expression (Figure 1C). This potentially suggests that the cell death signaling, observed upon co-expression of the *BnRPRs*, is activated from the cell nucleus, an aspect demonstrated for the RRS1/RPS4 pair upon effector recognition (Sarris et al., 2015). Apparently, and as it was previously hypothesized, (Qi and Innes, 2013), pathways other than the recently described Resistosome-related pathway, operate leading to cell death (Wang et al., 2019a; Wang et al., 2019b; Martin et al., 2020; Bi et al., 2021).

Regarding the mode of function of the *Bn*RPR pair, they seem to work in a fashion more similar to the Pik-1/Pik-2 pair (Zdrzalek et al., 2020) and not through negative regulation as documented for RRS1/RPS4 (Huh et al., 2017) or RGA4/RGA5 (Cesari et al., 2014b). In the *Bn*RPR pair, the presence of both NLRs is a prerequisite for cell death activation. While for the TNL pair RRS1/RPS4 and the CNL pair RGA4/RGA5, the executor NLR (RPS4 and RGA4, respectively), are able to activate cell death, however this is suppressed by its paired NLR, in the resting state (Cesari et al., 2014b; Maqbool et al., 2015; Huh et al., 2017). For the RRS1/RPS4 and RGA4/RGA5 pairs, it seems that intermolecular interactions act as a lock to limit costly defense activation and they can be circumvented through the rise of deleterious point mutations which destabilize the complex (Sohn et al., 2014; Newman et al., 2019) For the *Bn*RPRs pair, similar deleterious mutations in one member of the pair, would potentially lead to complex dissociation that will not however activate cell death. This adds up an extra level of NLRs regulation in order to avoid the unwanted effect of a random activation. On the other hand, this could also indicate that effectors can disrupt the complex formation, which could effectively block the immune response activation. A difference however, between the *Bn*RPRs and the Pik1/2 pair, relates to the autoactivation process. The *Bn*RPR pair, under the regulation of the native promoter or the 35S promoter, constitutively activates the cell death in *Nicotiana* species, in the absence of an elicitor. This cell death activation seems to be dependent on the two helper proteins tested here, EDS1 and NRG1, since, the cell death was abolished in the *eds1* and *nrg1* knockout mutants in *N. benthamiana* (Figure 2B).

However, this cell death activation cannot be attributed to the higher expression levels of *Bn*RPRs, due to the strong 35S promoter, since we observed that the use of the native *Bn*RPRs promoter also led to a conditional onset of cell death. In contrast, for the Pik1/2 pair that also seems to work in a way independent of the negative regulation, the overexpression by the use of a strong promoter (mas promoter) did not result in cell death activation (Zdrzalek et al., 2020).

Interestingly, the regulatory mechanism of *Bn*RPR pair seems to be more tightly regulated in Arabidopsis. When the *A. thaliana* transgenic lines, expressing the *BnRPRs* locus under the native promoter regulation, were grown under the same environmental conditions, no cell death induction was detected (Sup. Figure 5). At this point we cannot account for the exact pathway that is affected, since, these moderate environmental changes affect a number of pathways from the transcriptional to the post-translational level (Haak et al., 2017). This interesting finding needs to be further investigated in the future.

Regarding the role of the subcellular localization of the *Bn*RPR pair in the cell death activation process, we observed that changes on their localization, negatively affect the onset of cell death, a phenomenon frequently reported for NLRs (Qi and Innes, 2013). A shift from nuclear to cytoplasmic/nuclear localization results to a significant delay in the emergence of the cell death in the *Bn*RPR1 truncations lacking the TFSIIN domain (Figure 3C & Sup. Figure 3C). However, the deletion of the TFSIIN domain seems to be produced naturally as *in silico* predicted and also verified here experimentally (Sup. Figure 4B). Thus, alternative splicing (AS), as reported for other NLRs (Yang et al., 2014), seems to have a role on the correct activation process of the *Bn*RPR NLRs. At this point, it should be noted that, the TFSIIN domain contains one of the four predicted NLS residing in *Bn*RPR1 sequence. Thus, from the data presented here, we cannot deduce whether the lack of NLS or the TFSIIN domain *per se* is responsible for the observed changes in subcellular localization. However, both *Bn*RPR1 IDs are dispensable for cell death induction and probably, as shown for other IDs, have a role in effector recognition or affect developmental aspects (Grund et al., 2019).

In addition, and regarding the localization pattern of the examined proteins, we also observed a differential subcellular localization of *Bn*RPR2, when co-expressed with the truncated versions of *Bn*RPR1 lacking the TFSIIN domain. Similarly to *Bn*RPR1, *Bn*RPR2 could also be detected in the cytoplasm, which also indicates that *Bn*RPR2 is in complex with *Bn*RPR1.

Finally, we observed a co-regulation of the two *BnRPR*s genes. Upon co-expression, both *BnRPR1* and *BnRPR2* transcripts accumulate at lower levels than those observed when each gene is expressed alone, ultimately leading to lower protein accumulation levels (Figure 5C). The molecular mechanism for this phenomenon, is under investigation. RNA silencing mechanism could be a possible explanation, since, the two genes share complementary sequences at the 5’-ends of their transcripts that could potentially activate the production of nat-siRNAs, leading to the degradation of the transcripts and the downregulation at the transcriptional level. The only reference so far, related to a role of the nat-siRNAs on plant immunity concerns the nat-siRNAATGB2 (Katiyar-Agarwal et al., 2006). This nat-siRNAs is produced from the overlapping region of the 3’ untranslated regions (UTRs) of two mRNAs that are transcribed from adjacent genetic loci, Rab2-like small GTP-binding protein gene (*ATGB2*) and Pentatricopeptide repeats protein-like gene (*PPRL*) and is induced by the bacterial pathogen *Pseudomonas syringae* carrying effector *avrRpt2*.

Altogether, in this study, we report the identification of a new pair of NLR-IDs from the economically important species *B. napus*, which induce cell death only upon co-expression. Thus, this NLR-ID pair adopts a different mode of action from other paired NLRs, since, the presence of both NLRs is a prerequisite to trigger cell death induction. Their subcellular localization is important on the onset of the cell death induction, while regulatory mechanism at the post-transcriptional level have an impact on their activation process. Our work highlights mechanisms related to expression regulation of NLR-IDs and cell death activation in different plant species, while also denotes that initiation of plant immunity is also activated from the nucleus and not only through the cytoplasmic Resistosome activation pathway (Mermigka et al., 2020; Bi et al., 2021).

## EXPERIMENTAL PROCEDURES

### Plant material and growth conditions

*N. benthamiana, N. tabaccum* (var. Petit Gerard) and *A. thaliana* (Columbia ecotype), *B. napus* (var.ZS11) plants were grown under greenhouse conditions.

### Generation of *A. thaliana* transgenic lines

*A. thaliana* transgenic lines expressing the *BnRPRs* locus were generated with the floral dip method (Clough and Bent, 1998) using *A. tumefaciens* (GV3101) expressing the pICSL86955OD:*BnRPR* construct described above. Selection of the transgenic plants was conducted with the application of the herbicide Basta (Bayer) as described by Clough and Bent (1998).

### Agroinfiltration

For the agroinfiltration assays, all constructs were transformed into *Agrobacterium tumefaciens*, strain C58C1. The agrobacteria carrying the different constructs were grown in liquid LB medium supplemented with appropriate antibiotics for 24 hours. Cell were harvested by centrifugation, washed twice with an equal volume as the initial culture with 10mM MgCl_2_, and re-suspended in infiltration medium (10 mM MgCl_2_, 10 mM MES [pH 5.6]). For the HR assays, the final OD_600_ for each transformed agrobacterium was adjusted at 0.5 and the HR was assessed two to five days post infiltration. For agroinfitration assays for purposes other than HR tests (confocal microscopy, Co-IP assays) the final OD_600_ was adjusted at 0.2 in order to avoid tissue necrosis. All assays were performed in 5-6 weeks old plants.

### Plasmid Construction

The Golden Gate method (Engler and Marillonnet, 2014) was used for the construction of most expression constructs. The vectors and modules were provided in the Golden Gate cloning kit from The Sainsbury Laboratory, Norwich, United Kingdom (Engler and Marillonnet, 2014). Sequences were amplified by PCR using Phusion polymerase (NEB). Amplicons were separated on 1.0 -1.8 % agarose gel, and isolated from the gel with the gel extraction kit (Macherey-Nagel). The amplicons were A-tailed and cloned with T/A cloning into a Golden Gate compatible version of pBluescript II SK(-) developed in our lab. Cloning into the final expression vectors (pICH86988) was then performed by Dig-Lig reaction. In order to clone the whole genomic sequence of the *BnRPR* locus, PCR was conducted to amplify the whole sequence (Phusion polymerase, NEB), the amplicon was A-tailed and ligated into a T-tailed version of pICSL86955OD with T4 DNA ligase (NEB). The T-tailed version of pICSL86955OD was prepared according to (Zhou and Gomez-Sanchez, 2000).

### Genomic DNA extraction

For screening purposes in *A. thaliana* transgenic lines the protocol for fast extraction of genomic DNA was used (https://openwetware.org/wiki/Arabidopsis_gDNA_isolation). For high quality gDNA isolation DNeasy Plant Mini Kit (Qiagen) was used.

### RNA extraction and RT-PCR

Total RNA was isolated from deep-frozen plant material using the TRIzol^®^ method (Invitrogen) according to the manufacturer’s specifications. For cDNA synthesis 2 μg of DNaseI (Roche) treated total RNA was reverse transcribed using M-MuLV reverse transcriptase (Minotech). Rt-PCR was conducted with varying sets of primers (Supplementary Table 1).

### Real Time PCR

The expression analysis of *BnRPR1, BnRPR2* and the reference genes was determined using real-time PCR. Each cDNA sample was ten times diluted and amplified using KAPA SYBR^®^ FAST qPCR Kit (Kapa Biosystems) on the CFX Connect™ Real-Time PCR detection System (Bio-Rad). For real time PCR reaction, 2μl of ten times diluted cDNA was used as template in a reaction containing 1X of Kapa Master Mix, 0.2 μM primers in 10μl final volume. Each cycle consisted of denaturation at 95 °C for 3 sec and annealing at 60 or 62 °C (depending on the primer) for 20s. Ubiquitine and NPTII (Neomycin phosphotransferase) or actin were used as reference genes. All primers used in real time PCR are listed in Supplementary Table 1. The experiments were performed in three biological replicates. Analysis was conducted as described by (Pfaffl, 2001; Pfaffl et al., 2002).

### Co-IP assay and Immunodetection

Co-IP assays were performed as described by (Sarris et al., 2015). Briefly, 600mg of plant tissue was homogenized in 2 ml Co-IP buffer (10% (v/v) glycerol, 100 mM Tris (pH=8), 1 mM EDTA, 150 mM NaCl, 5 mM DTT, 0,2% Np-40 and protease inhibitors (Sigma). The lysate was centrifuged at 10.000 g for 20 min at 4 °C and a 500 μl aliquot was used for Co-IP assay. The lysates were incubated with 1.5 μl of monoclonal Anti-Flag M2 (Sigma) for 2 hrs at 4 °C with gentle agitation followed by the addition of 25 μl of Protein A/G magnetic beads (Pierce) and incubation for another 2 hour period. Three washes with Co-IP buffer were performed and the beads were resuspended in 50 μl o Co-IP buffer for further use. Proteins were separated in 8% SDS-PAGE mini-gel (Bio-Rad), blotted to PVDF membrane (Millipore) by wet-transfer for 90 min at 300 mA. For western analysis the membrane was incubated for 1 hr in blocking buffer (TBS-T + 5% milk), washed twice with TBS-T, incubated for 1 hr in TBS-T + 3% milk + α-FLAG (Sigma) or a-MYC (1:4,000 dilution) followed by three washes with TBS-T and 1 hr incubation in TBS-T + 1% milk + anti-mouse serum (1:10,000) (Promega). All steps were carried out at room temperature. Signals were detected using the SuperSignal West Pico Chemiluminescent Substrate (Thermo Scientific). Stripping of the membranes was performed by incubating the membrane for 30 minutes in stripping solution (200 mM glycine pH = 2.5, 1% SDS), followed by two washes in TBS-T for 15 minutes.

### Confocal microscopy

Sections of *N. benthamiana* agroinfiltrated leaves (2dpa) were analyzed with a Leica SP8 confocal microscope. YFP and mCherry were excited at 514 and 561, respectively. Emission for YFP and mCherry was recorded at 520-530 nm and 600-650 nm, respectively.

## Acknowledgements

The authors would like to thank Synthetic Biology team and Mr Mark Youles at The Sainsbury Laboratory for providing the Golden Gate cloning kit. The authors would like to thank Professor Jonathan DG Jones (The Sainsbury Laboratory), for providing the *N. benthamiana eds1* and *nrg1* knockout mutants, as well as, for the helpful comments on the manuscript.

## Authors’ contributions

P.F.S. and G.M. designed the research. G.M., A.A., A.M., and N.A. performed the research. P.F.S. and G.M. wrote the paper. All authors have read and approved the manuscript.

## Supplementary Figures and Tables

**Supplementary Table 1:** Primer sequences used in this study.

**Sup. Figure 1: Expression of *BnRPR1* and *BnRPR2* in the leaves of 10 day old *B. napus* plants**.

RT-PCR was performed on cDNA from the leaves of 10 day old *B. napus* plants with specific primers to amplify part of *BnRPR1* and *BnRPR2* while Actin was used as an internal control for the successful generation of cDNA. Expression of both transcripts was confirmed. Expected sizes: *BnRPR1* =189 bp, *BnRPR1* = 123 bp, *Actin* = 316 bp.

**Sup. Figure 2: Transient co-expression of *BnRPR1* and *BnRPR2* tagged with different epitopes triggers an autoimmune response**.

*N. tabacum* leaves were agroinfiltrated with the different constructs and the progression of HR was monitored. All BnRPR1/BnRPR2 combinations lead to cell death. The combination *Bn*RPR1-YFP/ *Bn*RPR2-MYC and *Bn*RPR1-nVENUS/ *Bn*RPR2-mCherry induced the fastest response while the combination *Bn*RPR1-nVENUS/ *Bn*RPR2-cCFP induced a delayed response.

**Sup. Figure 3: Subcellular localization and cell death induction of the three *Bn*RPR1 truncations**.

(a) Graphical representation of the different truncations of *Bn*RPR1, lacking the B3 domain (*Bn*RPR1ΔB3), the TFSIIN domain (*Bn*RPR1ΔTFSIIN) or both (*Bn*RPR1ΔB3ΔTFSIIN).

(b) Truncations of BnRPR1 are also identified in the cytoplasm. *N. benthamiana* leaves were agroinfiltrated with 35S:BnRPR1-YFP, 35S:*BnRPR1ΔB3-YFP, 35S:BnRPR1ΔTFSIIN-YFP* and *35S:BnRPR1ΔB3ΔTFSIIN-YFP* and analyzed with confocal microscopy two dpa. Truncations of *BnRPR1* have a nucleocytoplasmic localization pattern.

(c) *Bn*RPR1 truncated proteins lacking the TFSIIN domain lead to delayed cell death. *Bn*RPR2:mCherry was transiently co-expressed with *Bn*RPR1-YFP and the three truncations (*Bn*RPR1ΔB3-YFP, *Bn*RPR1ΔTFIIN-YFP and *Bn*RPR1ΔB3ΔTFIIN-YFP) in *N. benthamiana* leaves and cell death was monitored over a period of four days. Cell death was observed in the *Bn*RPR1 truncated proteins lacking the TFSIIN domain with a one day compared to the other combinations. Red dots denote the macroscopic cell death symptom.

**Sup. Figure 4: Characterization of transgenic *A. thaliana* plants expressing the *Bn*RPRs locus**.

(a) Comparison of the plant growth between *A. thaliana* w.t. and two Col^BNRPR^ T2 transgenic lines (Col^BNRPR^ l2 and Col^BNRPR^ l4) in different developmental stages (juvenile and adult stage, left and right panel, respectively). Col^BNRPR^ transgenic plants have similar growth and development to w.t. plants.

(b) RT-PCR was conducted on cDNA extracted from the Col^BNRPR^ T2 transgenic plants. Three siblings of each line were analyzed for the expression of *BnRPR1* and *BnRPR2*. In all cases both genes accumulate in the transgenic lines. In the case of *BnRPR1*, two bands are detected corresponding to the *in silico* predicted spliceforms. One splice form corresponds to the fully spliced sequence while the second splice form, contains a premature stop codon due to intron retention. Actin was used as an internal control for the successful generation of cDNA while the no-RT-PCR is also included. (-) and (+) correspond to negative (dH2O) and positive control (agroinfiltrated tissue with *BnRPR1* and *BnRPR2* constructs), respectively, that were included in the PCR. Expected sizes: *BnRPR1* = 271/ 195 bp, *BnRPR2* = 123 bp, *Actin* = 316 bp.

(c) Clustal W Alignment of part of the detected spliceforms of *BnRPR1* shown in (b). Spliceform 1 corresponds to the fully spliced sequence while the Spliceform 2 contains a premature stop codon due to intron retention and corresponds to the truncated isoform lacking the TFSIIN domain.

**Sup. Figure 5: A moderate change in the growth conditions did not induce cell death in the *A. thaliana* Col**^**BNRPR**^ **T2 transgenic lines**.

Comparison of the plant growth between *A. thaliana* w.t. and two Col^BNRPR^ T2 transgenic lines (Col^BNRPR^ l2 and Col^BNRPR^ l4 and Col^BNRPR^ l10) grown for a period of 14 days under different developmental conditions (Cond. I: 22 °C 50% RH vs Cond. II: 26 °C 80% RH, left and right panel, respectively). W.t. and Col^BNRPR^ transgenic plants show a stressed phenotype under Condition II.

## Graphical Abstract

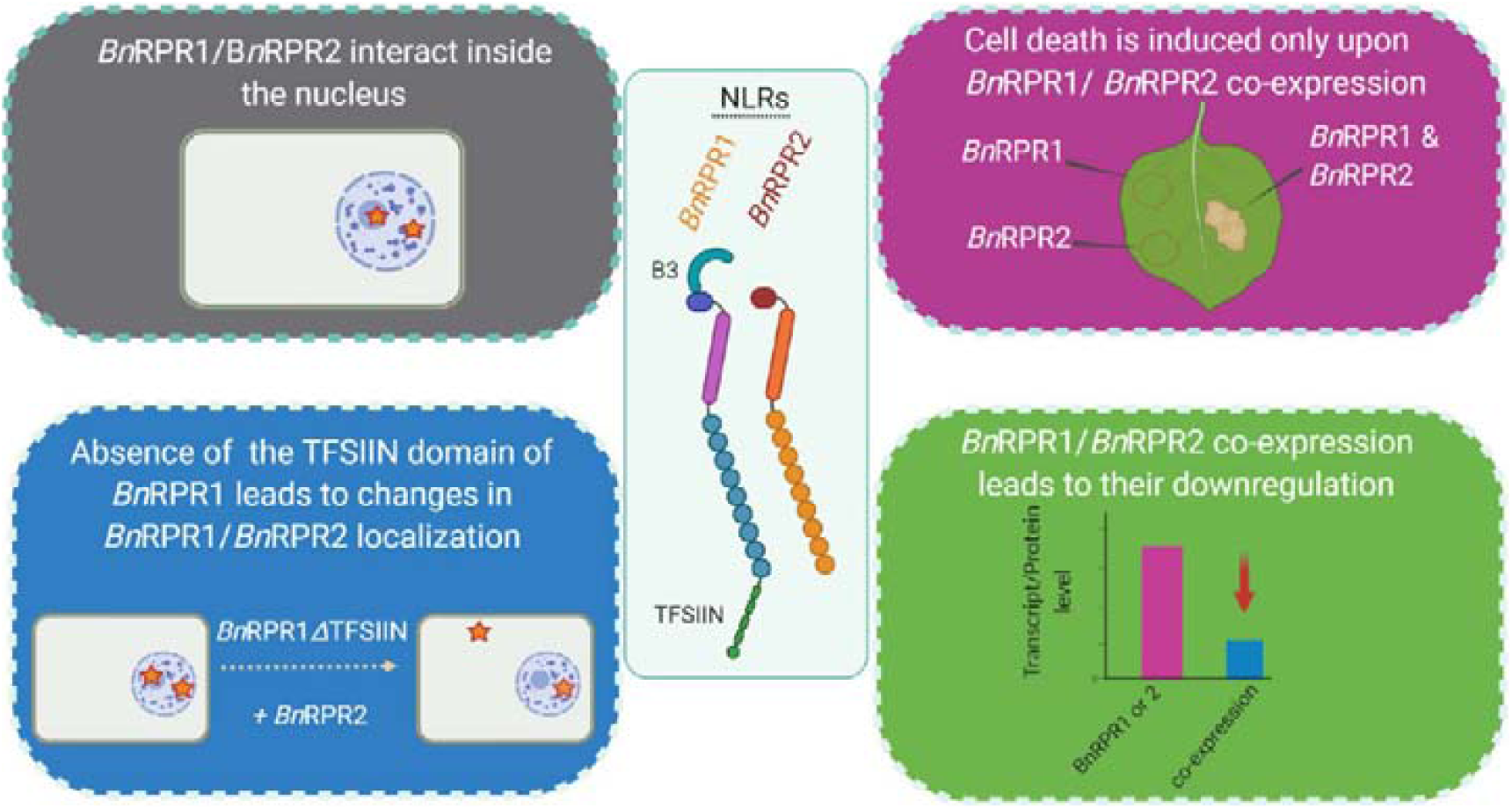

