## Supplementary Figures for "Assassination Tango: An NLR/NLR-ID immune receptors pair of rapeseed co-operates inside the nucleus to activate cell death"

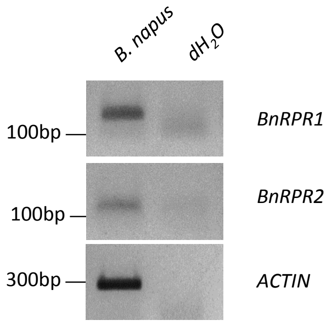


**Sup. Figure 1: Expression of *BnRPR1* and *BnRPR2* in the leaves of 10 day old *B. napus* plants.**

RT-PCR was performed on cDNA from the leaves of 10 day old *B. napus* plants with specific primers to amplify part of *BnRPR1* and *BnRPR2* while Actin was used as an internal control for the successful generation of cDNA. Expression of both transcripts was confirmed. Expected sizes: *BnRPR1* =189 bp, *BnRPR1* = 123 bp, *Actin* = 316 bp,

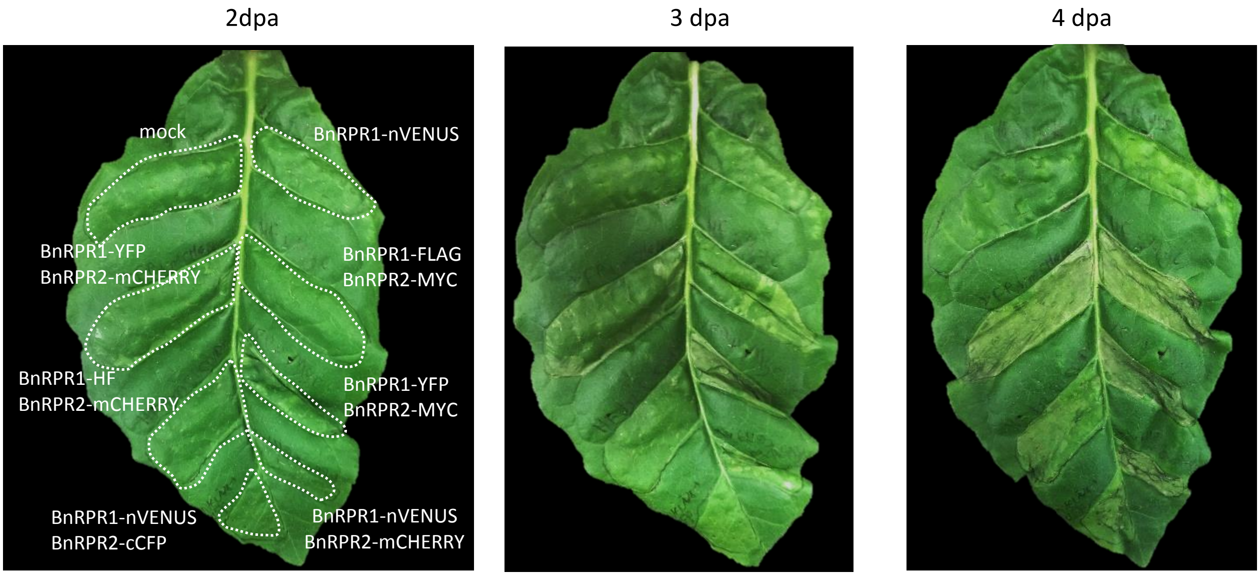


**Sup. Figure 2:** Transient co-expression of *BnRPR1* and *BnRPR2* tagged with different epitopes triggers an autoimmune response. *N. tabacum l*eaves were agroinfiltrated with the different constructs and the progression of HR was monitored. All BnRPR1/BnRPR2 combinations lead to cell death. The combination *Bn*RPR1-YFP/ *Bn*RPR2-MYC and *Bn*RPR1-nVENUS/ *Bn*RPR2-mCHERRY induced the fastest response while the combination *Bn*RPR1-nVENUS/ *Bn*RPR2-cCFP induced a delayed response.

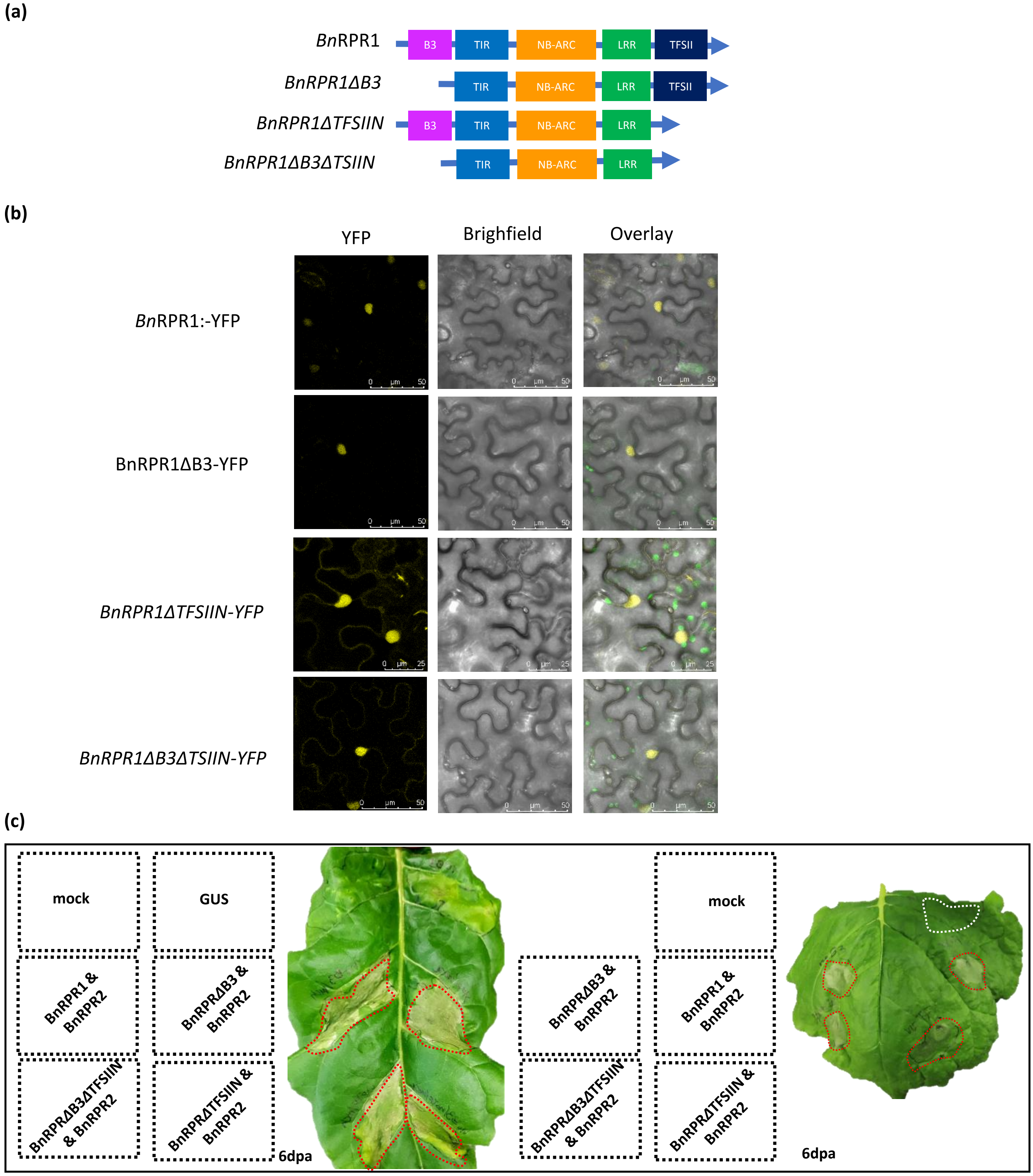


**Sup. Fig. 3: Subcellular localization and cell death induction of the three *Bn*RPR1 truncations.**

(a) Graphical representation of the different truncations of *Bn*RPR1, lacking the B3 domain (*Bn*RPR1ΔB3), the TFSIIN domain (*Bn*RPR1ΔTFSIIN) or both (*Bn*RPR1ΔB3ΔTFSIIN).

(b) Truncations of BnRPR1 are also identified in the cytoplasm. *N. benthamiana* leaves were agroinfiltrated with *35S:BnRPR1-YFP*, *35S:BnRPR1ΔB3-YFP*, *35S:BnRPR1ΔTFSIIN-YFP* and *35S:BnRPR1ΔB3ΔTFSIIN-YFP* and analyzed with confocal microscopy two dpa. Truncations of *BnRPR1* have a nucleocytoplasmic localization pattern.

*(c)* *Bn*RPR1 truncated proteins lacking the TFSIIN domain lead to delayed cell death. *Bn*RPR2:mCHERRY was transiently co-expressed with *Bn*RPR1-YFP and the three truncations (*Bn*RPR1ΔB3-YFP, *Bn*RPR1ΔTFIIN-YFP and *Bn*RPR1ΔB3ΔTFIIN-YFP) in *N. benthamiana* leaves and cell death was monitored over a period of six days. Cell death was observed in the *Bn*RPR1 truncated proteins lacking the TFSIIN domain with a two to three day compared to the other combinations. Red dots denote the macroscopic cell death symptom.

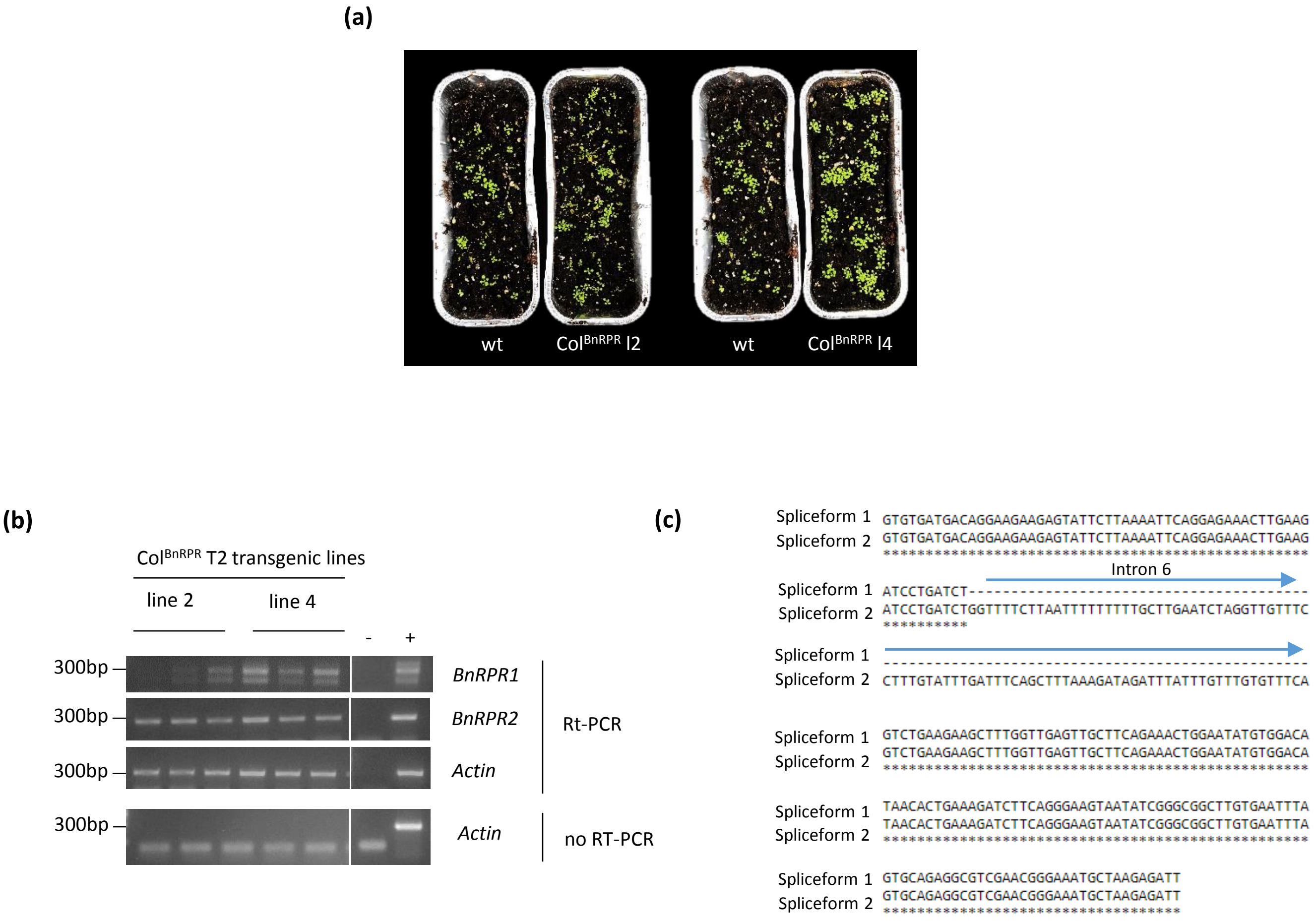


**Sup. Figure 4: Characterization of transgenic *A. thaliana* plants expressing the *BnRPRs* locus.**

(a) Comparison of the plant growth between *A. thaliana* w.t. and two Col$^{BnRPR}$ T2 transgenic lines (Col$^{BnRPR}$ l2 and Col$^{BnRPR}$ l4) in different developmental stages (juvenile and adult stage, left and right panel, respectively). Col$^{BnRPR}$ transgenic plants have similar growth and development to wt plants.

(b) Rt-PCR was conducted on cDNA extracted from the Col$^{BnRPR}$ T2 transgenic plants. Three siblings of each line were analyzed for the expression of *BnRPR1* and *BnRPR2.* In all cases both genes accumulate in the transgenic lines. In the case of *BnRPR1,* two bands are detected corresponding to the *in silico* predicted spliceforms. Actin was used as an internal control for the successful generation of cDNA while the no-RT-PCR is also included. (-) and (+) correspond to negative (dH2O) and positive control (agroinfiltrated tissue with *BnRPR1* and *BnRPR2* constructs), respectively, that were included in the PCR. Expected sizes: *BnRPR1* = 271/ 195 bp, *BnRPR2* = 123 bp, *Actin* = 316 bp.

(c) Clustal-W Alignment of part of the detected spliceforms of *BnRPR1* shown in (b). Spliceform 1 corresponds to the fully spliced sequence while the Spliceform 2 contains a premature stop codon due to intron retention and corresponds to the truncated isoform lacking the TFSIIN domain.

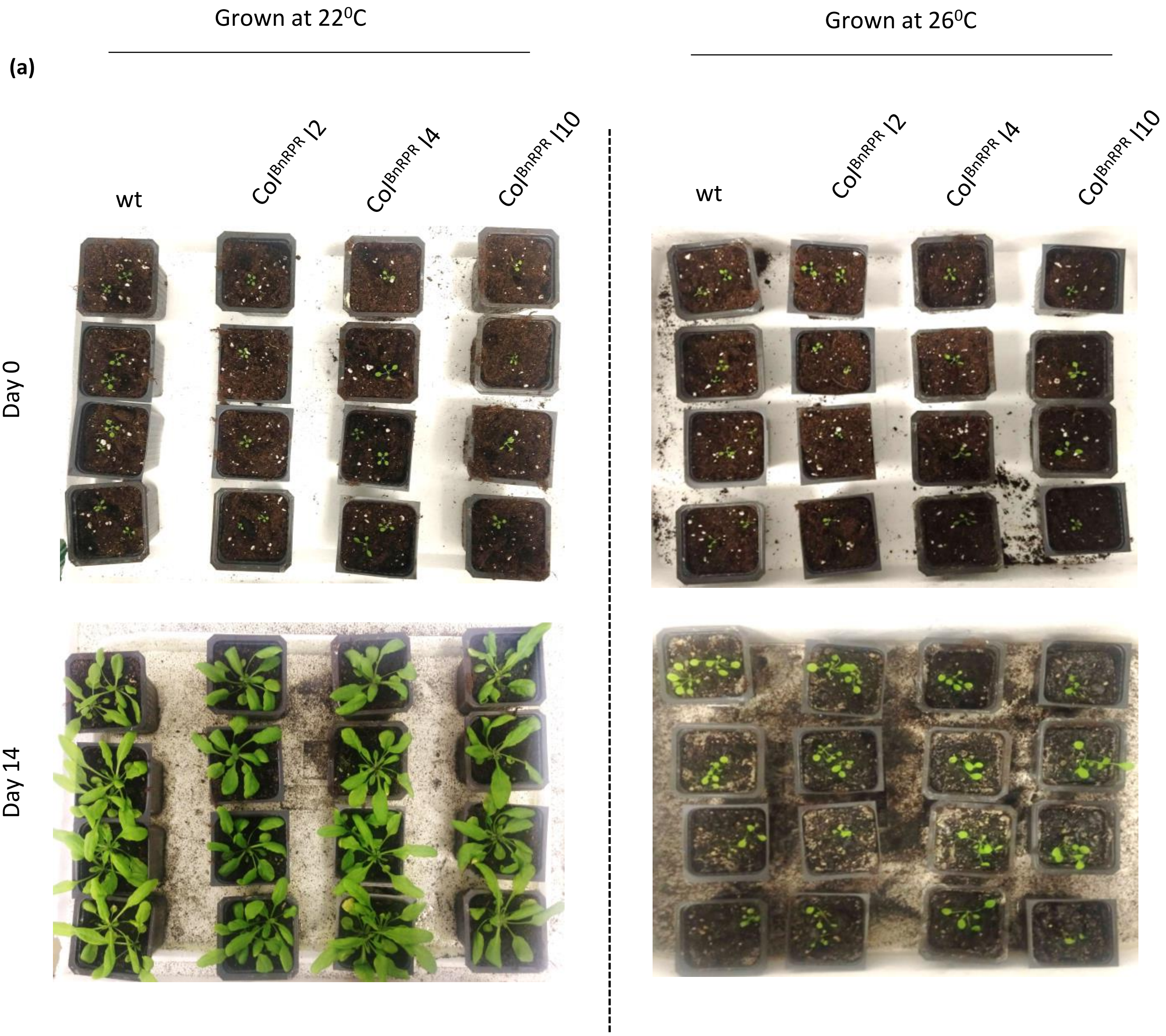


**Sup. Figure 5: A moderate change in the growth conditions did not induce cell death in the *A. thaliana* Col$^{BnRPR}$ T2 transgenic lines.** Comparison of the plant growth between *A. thaliana* w.t. and two Col$^{BnRPR}$ T2 transgenic lines (Col$^{BnRPR}$ l2 and Col$^{BnRPR}$ l4 an Col$^{BnRPR}$ l10) grown for a period of 14 days under different developmental conditions (Cond. I: $22^0$C 50% RH *vs* Cond. II: $26^0$C 80% RH, left and right panel, respectively). W.t. and Col$^{BnRPR}$ transgenic plants show a stressed phenotype under Condition II.
