## Supplementary Table for "Assassination Tango: An NLR/NLR-ID immune receptors pair of rapeseed co-operates inside the nucleus to activate cell death"

**Supplementary Table 1**: Primer sequences used in this study.

| a/a | Name | sequence |
| --- | --- | --- |
| 1 | BnPR1-Fw-1 | ΤTTGGTCTCAAATGCGGCAGCCATGGAAG |
| 2 | BnPR1-Rev-1 | TTTGGTCTCATGACGTGCTTCCCTTCCAAT |
| 3 | BnPR1-Fw-2 | TTTGGTCTCAGTCATTGGTGTCGGAGCTG |
| 4 | BnPR1-Rev-2 | TTTGGTCTCACTCATCAATGAGCCCATAGATTCCTA |
| 5 | BnPR1-Fw-3 | TTTGGTCTCATGAGTCTCTCATCAGCATTGTAG |
| 6 | BnPR1-Rev-3 | TTTGGTCTCAACCGATTGATGATGGCATTTC |
| 7 | BnPR1-Fw-4 | TTTGGTCTCACGGTGGGCTCAGTAAACTTGTTAC |
| 8 | BnPR1-Rev-4 | TTTGGTCTCACGAAGGTAGATGCAACCAAAACTGAGAGTC |
| 9 | BnPR1-Rev-4 Stop | TTGGTCTCAAAGCTCATAGATGCAACCAAAACTGAGAGTC |
| 10 | BnRPR1-B3-Fw | ΤTTGGTCTCAAATGGCTGTTCTCGGACAGTGTCAGG |
| 11 | BnRPR1-TFSII-Rev | TTTGGTCTCACGAAGGCCTGTCATCACACAAAGTAGCAA |
| 12 | BnPR2-Fw-1 | ΤTTGGTCTCAAATGGCTGCCGCATCTTCC |
| 13 | BnPR2-Rev-1 | TTTGGTCTCACCGAGGGCTCAACCAAATATAAACC |
| 14 | BnPR2-Fw-2 | TTTGGTCTCATCGGTTTACCTTTTTAAAGGAGGTTG |
| 15 | BnPR2-Rev-2 | TTTGGTCTCACGAAGGCCAGATATTAGGCTTTGCTGGC |
| 16 | BnPR2-Rev-2 stop | TTTGGTCTCAAAGCTCACCAGATATTAGGCTTTGCTGGC |
| 17 | BnRPR1-Fw splice | GTGTGATGACAGGAAGAAGAGTA |
| 18 | Bn-1 Rev splice | AATCTCTTAGCATTTCCCGTTCG |
| 19 | qBnRPR1-Fw | CAGACTTCCATATAGTGTGTCAAACG |
| 20 | qBnRPR1-Rev | GAGACCAATGACGTGCTTCC |
| 21 | qBnRPR2-2 Fw | GTGCTGAAAGTGTGCTATGAGG |
| 22 | qBnRPR2-Rev | GGTATGGTGAGTGCTCAGAAC |
| 23 | qNPTII-Fw | CTGCCGAGAAAGTATCCATCA |
| 24 | qNPTII-Rev | GATGTTTCGCTTGGTGGTCG |
| 25 | qUBI3-Fw | GCCGACTACAACATCCAGAAGG |
| 26 | qUBI3-Rev | TGCAACACAGCGAGCTTAACC |
| 27 | qACT-Fw | GGAGATGATGCTCCAAGAGC |
| 28 | qACT-Rev | CGATTAGCCTTTGGGTTAAGAGG |
